# The nitrogenase G-subunit is an ancient orphan protein that drove the ecological expansion of nitrogen fixation

**DOI:** 10.1101/2023.04.03.535378

**Authors:** Bruno Cuevas-Zuviría, Amanda K. Garcia, Brooke M. Carruthers, Betül Kaçar

## Abstract

Nitrogenase metalloenzymes have catalyzed biological nitrogen fixation for billions of years and revolutionized planet Earth by supplying essential nitrogen to the biosphere. How these enzymes were built and distributed by microbial and evolutionary processes in a shifting geochemical landscape remains an open question. Here, we probe the birth and evolution of the G-subunit protein, an integral, Precambrian-age structural component of certain nitrogenase isozymes that makes its appearance midway through nitrogenase evolutionary history. We establish that the G-subunit is an orphan protein, with no homologs detected across wider protein diversity. We find that G-subunit emergence accompanied both the diversification of nitrogenase metal usage and an ecological expansion of nitrogen-fixing microbes during the transition in enviromental metal availabilities triggered by Earth surface oxygenation ∼2.5 billion years ago. Further, analyses of ancestral nitrogenase structures implicate a role for the G-subunit in novel metal incorporation, which would have primed nitrogenases and their hosts to exploit these newly diversified geochemical environments. However, permanent recruitment of the G-subunit into the nitrogenase complex was likely only enabled by tuning preexisting, protein interaction features that were selected prior to Earth oxygenation. Our results showcase how contingent evolutionary novelties shape microbial ecological responses and their global consequences.

## INTRODUCTION

Over billions of years, life generated an enormous wealth of biomolecular diversity, producing an estimated 10_5_ extant protein families^1^ and a much larger multitude of extinct proteins. Through this process, biology became a defining component of the Earth system, which experienced tremendous revolutions in climate and biogeochemistry that are largely attributable to the molecular innovations of life itself^2^. A unified understanding of the Earth-life system requires investigating how the proteins and biogeochemical processes that power planetary phenomena emerged, evolved and proliferated.

In many cases, nature was a tinkerer^3^, generating novel functions through the progressive duplication, accretion, and recombination of pre-existing protein elements^4-7^. In rarer cases, complete novelty arose through the *de novo* birth of genes^4,8^. This scenario represents a more difficult evolutionary hurdle due to the improbability that formerly noncoding DNA would produce a structurally or functionally coherent peptide^9^. Retracing these different evolutionary events in the histories of biogeochemically critical enzymes is a considerable challenge given their antiquity, since many of these enzymes, including those required for essential elemental cycling, likely evolved more than three billion years ago^2,10,11^.

Here, we identify a case of molecular novelty in the evolutionary history of biological nitrogen fixation. Nitrogenase metalloenzymes provide the sole biochemical gateway for essential nitrogen into Earth’s biosphere, consequently transforming the planet^10,12-14^. They have done so for more than 3 billion years^12,13^. This metalloenzyme family is today represented by three homologous isozymes—molybdenum-(Nif), vanadium-(Vnf), and iron-only (Anf) nitrogenases, the latter two which evolved more recently from Nif^15^. Each are named for the differing compositions of their active-site metalloclusters but otherwise share a core, two-component oxidoreductase architecture^16-18^. The antiquity and biological significance of this architecture is highlighted by its conservation in distantly related enzymes within the nitrogenase superfamily^19^, the origin of which likely dates back to the last universal common ancestor^19,20^.

A striking distinction in all known Vnf and Anf nitrogenases is a specific protein subunit called the “G-subunit”^21-24^. G-subunit does not exist in any known Nif nitrogenases, and it is an essential component for nitrogen fixation in modern Vnf and Anf nitrogenases^23,25^. The origin and evolutionary role of the G-subunit is not known. We know that the adaptive landscape of nitrogenases shifted with transitions in environmental metal availabilities brought on by Earth surface oxygenation ∼2.5 billion years ago^26,27^ which may have impacted nitrogen fixation diversification^15,26,28^. To what degree was a novelty such as the G-subunit necessary for the expansion of metal usage in nitrogen fixation?

In the present study, we elucidate the evolutionary history of the nitrogenase G-subunit and address the specific interplay between molecular evolutionary events and environmental triggers in the ecological diversification of biological nitrogen for the first time. We infer the ancestral sequences and structures of nitrogenase enzymes through the emergence of the G-subunit, replaying the sequence of events that gave rise to this nitrogenase protein component >1.5 billion years ago. We further interrogate the genetic and environmental context of its emergence, recruitment, and functional significane within the context of an ancient ecosystem.

## RESULTS

### The nitrogenase G-subunit is an ancient orphan protein

In modern nitrogen-fixing microbes, the nitrogenase G-subunit protein is a small, structural element of the larger, two-component enzyme complex. The first component is a H-subunit homodimer that delivers electrons to the second, catalytic component, which, in Nif, is a D- and K-subunit heterotetramer but, in Vnf/Anf (collectively referred to as “alternative nitrogenases”), is a D-, G-, and K-subunit heterohexamer^21^ (**Figure 1A**). Thus, the G-subunit is a primary distinction in the otherwise structurally comparable Nif, Vnf, and Anf complexes. To date, the sole, publicly available crystallographic structure of the G-subunit is from *A. vinelandii* Vnf^21^, a 113-amino-acid, four-helix protein (**Figure 1B**; the *A. vinelandii* Anf crystallographic structure has only recently been described^29^). In both Vnf and Anf, the G-subunit neighbors the D-subunit that houses the active-site metallocluster where N_2_ is bound and reduced^30^. The metal content of the metallocluster differs between Nif, Vnf, and Anf nitrogenase isozymes, incorporating Fe and Mo (“FeMo-co”), Fe and V (“FeV-co”), or only Fe (“FeFe-co”), respectively. Finally, though sharing a common ancestry^15,31,32^ and core N_2_-reduction mechanism^33^, Nif, Vnf, and Anf nitrogenases vary in their kinetic behavior and reactivities to different substrates^18,33-35^.

**Figure 1.**
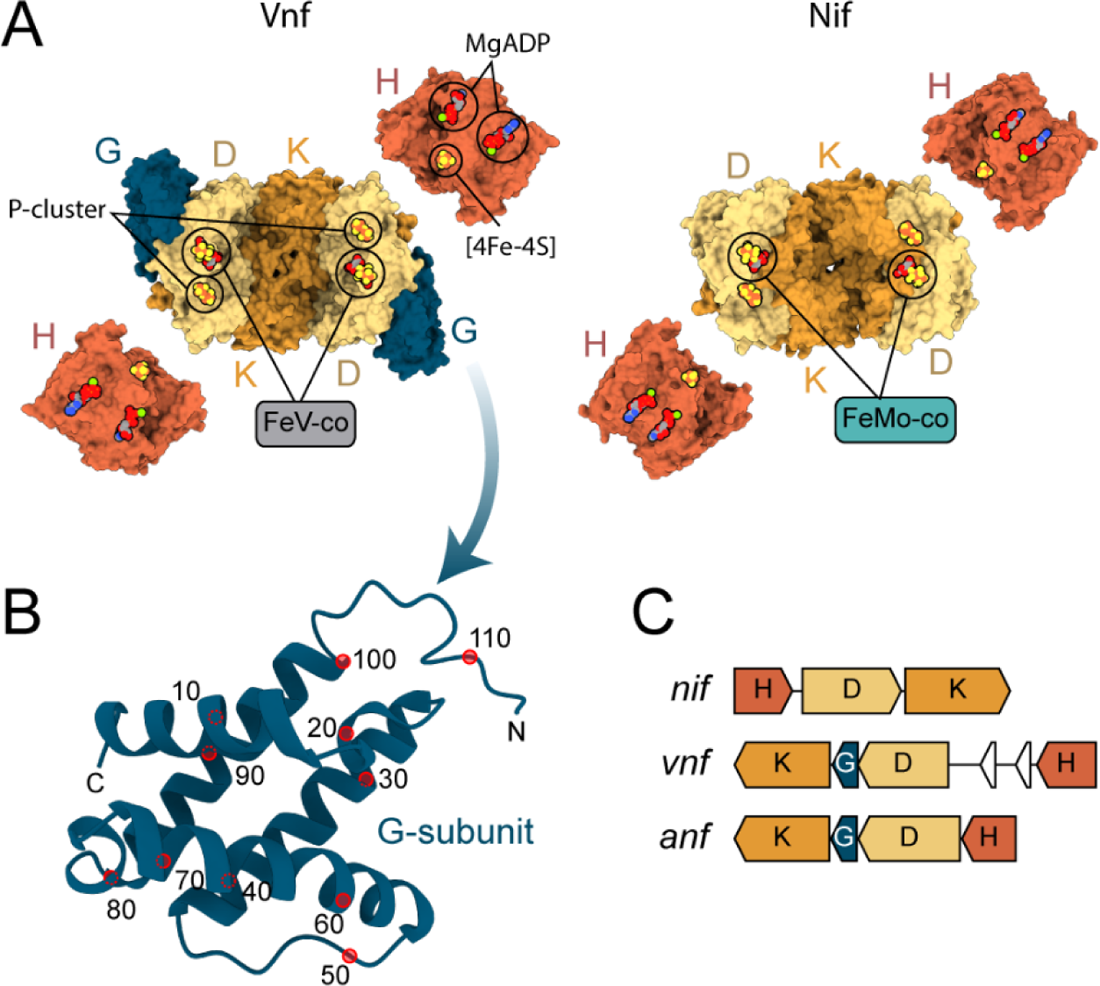
Structure and genetics of the nitrogenase G-subunit. **A)** Left, crystallographic structure of the extant A. vinelandii Vnf nitrogenase complex (VnfH, PDB 6Q93^36^; VnfDGK, PDB 5N6Y^21^) containing the G-subunit (dark blue). The Anf nitrogenase is inferred to be comparable in its multimeric structure, binding FeFe-co instead of FeV-co. Right, crystallographic structure of the extant Nif nitrogenase complex (NifH, PDB 1M34^37^; NifDK, PDB 3U7Q^38^) lacking the G-subunit. **B)** Secondary structure of the Vnf G-subunit. **C)** Arrangement of the Nif, Vnf, and Anf structural genes in A. vinelandii. Vnf/AnfG (dark blue) is located between Vnf/AnfD and Vnf/AnfK genes.

We began our evolutionary study of the nitrogenase G-subunit by constraining its place within the broader diversity of protein sequence and structure. Specifically, we sought to identify distant homologs that might provide insights into G-subunit ancestry and function. Though genetic studies have established the G-subunit as essential for an active, N_2_-reducing Vnf/Anf nitrogenase complex^23,25^, its precise functional contribution remains enigmatic. We implemented a homology search workflow that included both sequence-based (BLASTP, TBLASTN^39^, PSI-BLAST^40^, HMMER^41^) and structure-based (Foldseek^42^) methods appropriate for detection of distantly related proteins. Our search returned >400 canonical G-subunit proteins, but, notably, we did not find convincing similarity to any other protein. PSI-BLAST and HMMER searches together returned <10 hits that we could exclude from the canonical G-subunit protein family (i.e., were not encoded by genes within *vnf*/*anf* structural operons) (**Table 1, Figure S1**). In both cases, only short sequence regions of scattered taxonomic affinities aligned to our query with poor statistical significance. We were thus unable to rule out spurious matches. Additionally, we used the structure-based method, FoldSeek, leveraging the greater conservation of protein structure compared to sequence^43^ in homology detection. We identified two uncharacterized, AlphaFold-predicted structures that are not canonical G-subunit proteins but nevertheless share its four-helix structural arrangement (**Table 1, Figure S2, S3**). However, sequence-level similarity to G-subunit proteins is negligible and, given the lack of functional annotations for these hits, their relevance to early G-subunit evolution is unclear. Finally, we note that we did not detect sequence or structural similarity to NafY and CowN proteins, both of which have previously been proposed to resemble the G-subunit in their interactions with the nitrogenase complex^21,44,45^.

**Table 1.** Summary of G-subunit homolog search results. *Template modeling score for structural alignments.

| Method | Database | Total hits<br>(including<br>canonical G-subunit<br>proteins) | Distant<br>hits | E-value<br>range of<br>distant<br>hits | TM-score* of<br>distant<br>hits | Annotation of distant<br>hits | Taxonomy of<br>distant hits |
| --- | --- | --- | --- | --- | --- | --- | --- |
| BLASTP | NCBI non-redundant<br>proteins (nr) | 433 | 0 | n/a | n/a | n/a | n/a |
| TBLASTN | NCBI nucleotide<br>collection (nr/nt) | 205 | 0 | n/a | n/a | n/a | n/a |
| PSI-BLAST | NCBI non-redundant<br>proteins (nr) | 446 | 7 | 0.026-0.064 | n/a | TIGR04141 family<br>sporadically distributed<br>protein, Uncharacterized<br>protein | *Pseudomonas*<br>spp. |
| HMMER | UniProt<br>Reference<br>Proteomes | 94 | 1 | 0.061 | n/a | Uncharacterized protein | *Coregonus* sp. |
| FoldSeek | Protein Data<br>Bank | 3 | 0 | n/a | n/a | n/a | n/a |
| FoldSeek | AlphaFold-Uniprot/Swiss-Prot | 39 | 2 | 5.2-8.7 | 0.52-0.54 | Uncharacterized protein | *Vibrio vulnificus*,<br>*Deinococcus*<br>*grandis* |

If the G-subunit shares an ancient ancestry with another extant protein family, no strong vestige of this evolutionary relationship appears presently detectable. The absence of distant homologs to the G-subunit family is comparable to the case of “orphan genes,” which are individual genes that either have no resemblance to other sequences or have few, taxonomically restricted relatives^8,46^. Proposed mechanisms for the emergence of orphan genes include duplication followed by rapid divergence and *de novo* birth from previously non-genic DNA^4,8^. The latter has more recently been recognized as a significant source of orphan genes^47-49^ and thus would plausibly explain G-subunit origins. Regardless of the specific originating mechanism, our results indicate that the G-subunit family is a molecular innovation as unique to nitrogenases as it is to broader protein diversity.

### Invention of the G-subunit coincided with diversification of nitrogenase metal usage and host microbial ecology

We reconstructed phylogenies of nitrogenase G-subunit proteins and examined their genetic distribution across extant genomes and microbial host environments. We identified G-subunit homologs exclusively from *vnf* and *anf* gene clusters in bacterial and archaeal genomes (similarly, all identified *vnf*/*anf* clusters include the G-subunit), confirming that the G-subunit is a significant molecular distinction between Vnf/Anf and Nif nitrogenases^50^. In all analyzed clusters, *vnf/anfG* genes are located between *vnf/anfD and vnf/anfK* loci (**Figure 1C**). By contrast, *nifD* and *nifK* genes in our dataset are universally contiguous with small intergenic distances (median ≈ 25 bp) (**Figure S4**). Finally, all complete genomes harboring *vnf* and/or *anf* nitrogenase genes in our dataset also possess *nif* genes. Therefore, the presence of the G-subunit correlates with the capacity of a host diazotroph to express at least two different nitrogenase isozymes with distinct metal dependencies. We built two maximum-likelihood G-subunit phylogenies: the first from individual G-subunit protein sequences and the second from G-subunit sequences concatenated with other nitrogenase subunit sequences. Both trees show separate clustering of Vnf and Anf sequences, though deep branch support in the concatenated tree is significantly improved (**Figure 2, S5, S6**).

**Figure 2.**
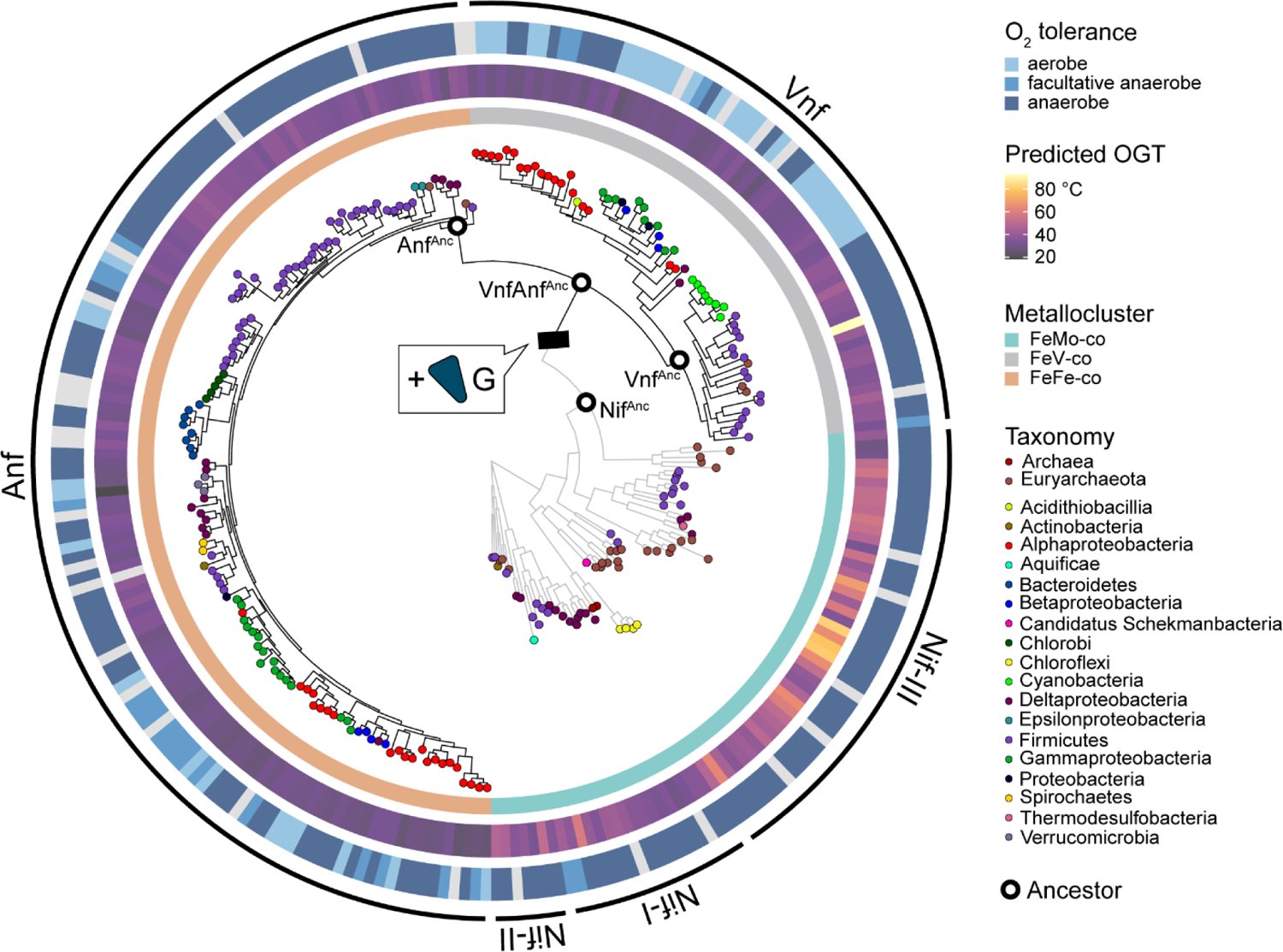
Evolutionary and environmental history of the nitrogenases and the G-subunit. Maximum-likelihood phylogeny built from concatenated nitrogenase subunit proteins. Oxygen tolerance, predicted optimal growth temperatures (OGT), and taxonomy of represented host microbes, as well as the metal content of the active-site metallocluster, are mapped to the phylogeny (light grey bars for O_2_ tolerance and predicted OGT indicate missing data). Nif clades are labeled according to the nomenclature used by Raymond et al.^31^ (e.g., Nif-I = Group I Nif, etc.). The G-subunit origin event is indicated by a black hash mark. Branches that track the emergence of the G-subunit and its diversification into Vnf and Anf groups are shown in black, whereas branches lacking the G-subunit are shown in grey. Ancestral nodes (Nif^Anc^, VnfAnf^Anc^, Vnf^Anc^, Anf^Anc^) targeted for sequence reconstruction and analysis are highlighted by bold outlined, white circles.

The overall topology of our concatenated nitrogenase tree mirrors previous phylogenetic analyses of nitrogenase proteins reconstructed without G-subunit sequences. Nesting of Vnf and Anf clades within Nif sequences indicates that Vnf/Anf are more recently evolved^15,32^, which, by extension, establishes the G-subunit as the most recently evolved structural component of the nitrogenase complex. Minimum age estimates of the Vnf/Anf clade range from ∼1.5 to 2.5 Ga, based on the timing of horizontal gene transfer events that pervade nitrogenase evolutionary history^51^. These estimates indicate that the Precambrian origin of the G-subunit roughly coincided with the initial accumulation of oxygen in the Earth surface environment^27^, the earliest whiffs of which have been identified from geochemical signatures in ∼2.5 to 3.0 Ga sediments^52,53^ and did not reach modern levels until ∼0.5 Ga^54^.

Our analyses demonstrate that the origin of the G-subunit was associated both with the gain of novel metal dependence and the ecological expansion of nitrogen fixation following the rise of atmospheric oxygen. Vnf and Anf sequences specifically branch from within the “Group III” nitrogenase lineage^31^, which is additionally populated by Nif homologs of either experimentally characterized or computationally predicted Mo-dependence^15,55^. Group III Nif are exclusively hosted by anaerobic taxa (primarily methanogenic Euryarchaeota and Firmicutes), many of which are probable thermophiles or hyperthermophiles (optimal growth temperatures predicted from genome content, see **Materials and Methods**) (**Figure 2, Supplementary Tables**). By contrast, Vnf/Anf are hosted by comparatively more taxonomically diverse and mesophilic archaea and bacteria that also harbor “Group I and II” Nif nitrogenases^31^, including the well-studied diazotrophic models like *A. vinelandii*, *Clostridium pasteurianum,* and *Rhodopseudomonas palustris*. Vnf/Anf are not hosted by organisms that harbor Group III Nif, despite Vnf/Anf emerging from within the Group III lineage. Within Group III, only Vnf and Anf clades contain sequences from aerobic taxa, including cyanobacteria which appear to only host Nif and Vnf nitrogenases. Together, these data indicate that G-subunit-containing Vnf and Anf nitrogenases were able to proliferate into ecological niches inaccessible to earlier diverged Group III Nif.

### An ancient assembly pathway preconditioned early nitrogenases for G-subunit interaction

To replay the first evolutionary steps in the birth of the G-subunit, we reconstructed and analyzed ancestral nitrogenase protein sequences. We examined the degree to which ancestral protein surface features were remodeled to accommodate a new subunit, with the expectation that G-subunit interactions would have driven the evolution of complementary residues within the interface. Because the G-subunit is exclusive to Vnf and Anf nitrogenases, the most parsimonious evolutionary scenario is that it emerged along the branch leading to the Vnf/Anf last common ancestor, VnfAnf^Anc^ (**Figure 2**). We therefore compared sequences and structures of nitrogenase ancestors reconstructed both immediately before (Nif^Anc^) and after (VnfAnf^Anc^) gain of the G-subunit.

We mapped the interface between ancestral G- and D-subunit proteins (“G-D interface”) by building its residue interaction network from the predicted VnfAnf^Anc^ structure (**Figure 3A-3D**). Nearly all well-conserved, G-subunit residues lie within the G-D interface and contribute significantly to its stability (**Figure S7**). The few exceptions are conserved residues distal from the interface with side chains oriented towards the interior of the helical bundle, which likely stabilize its globular arrangement.

**Figure 3.**
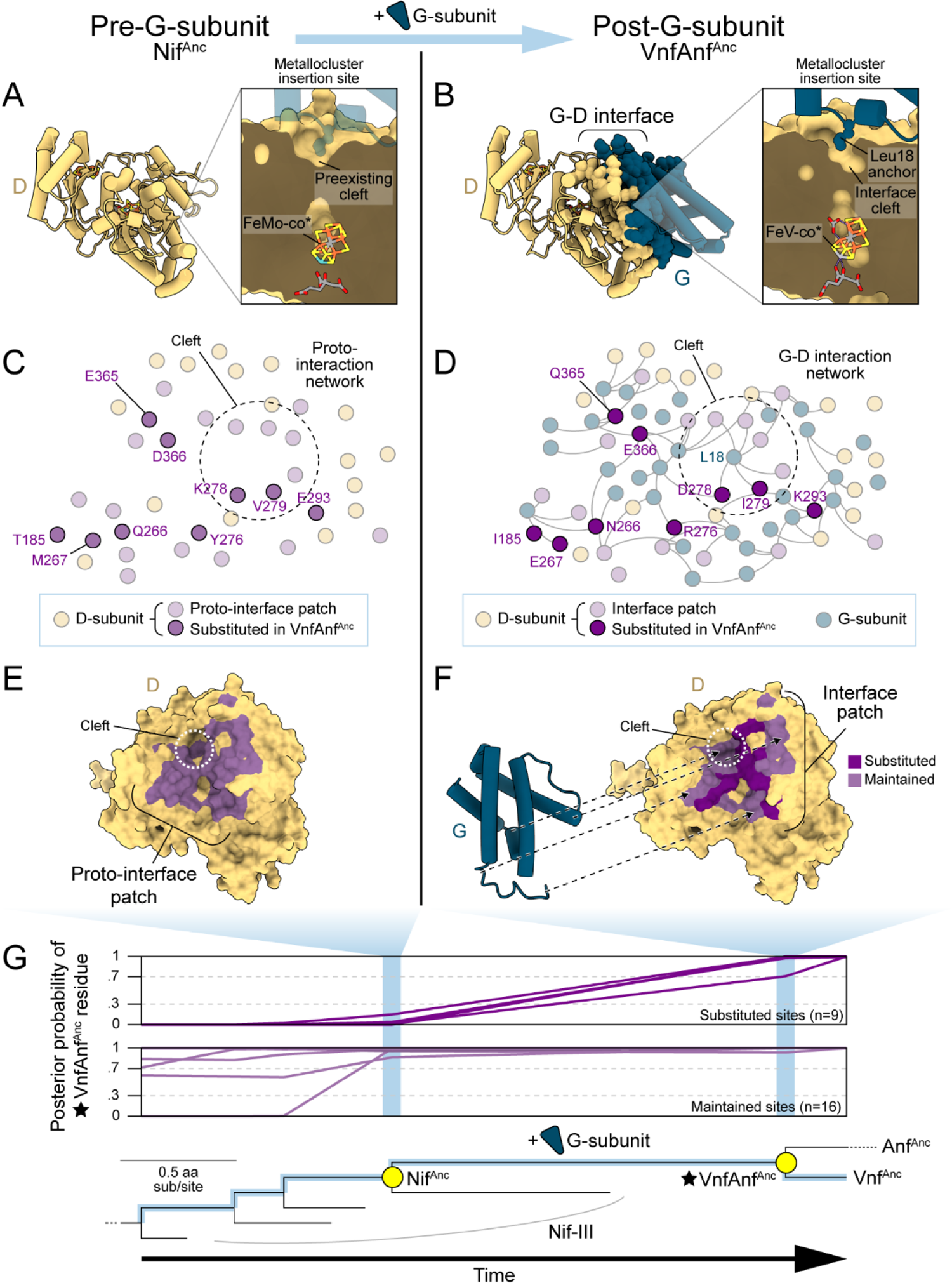
Birth of the nitrogenase G-subunit interface. **A)** Predicted Nif^Anc^ D-subunit structure, prior to G-subunit emergence. Inset shows cross-section at the putative metallocluster insertion site, the entrance of which forms a “cleft” in the D-subunit surface. Superimposed G-subunit structure (transparent) highlights relative position of the cleft. **B)** Predicted VnfAnf^Anc^ D- and G-subunit structures, following G-subunit emergence, shown in the same view as A). Side chain of the G-subunit Leu18 residue is anchored within the D-subunit cleft. **A-B)** Heterometal content for the ancestor metalloclusters is inferred. **C)** Nif^Anc^ G-D proto-interaction network. Purple residues are located within an interface patch conserved in extant Vnf/Anf. Residues that are later substituted in VnfAnf^Anc^ are shown bold and labeled to permit comparison with D). Residues within the cleft are circled. **D)** VnfAnf^Anc^ G-D interaction network, shown in the same view as C). Residues newly substituted in VnfAnf^Anc^ are shown in dark purple and labeled. **E)** Surface view of the Nif^Anc^ D-subunit proto-interface patch (purple). **F)** Surface view of the VnfAnf^Anc^ D-subunit interface patch, shown in the same view as E). Residues that are maintained between NifAnc and VnfAnfAnc are shown in purple, and those that are substituted are shown in dark purple. **G)** Evolution of residues within the interaction patch through the emergence of the G-subunit. For substituted sites, the posterior probability of the most likely VnfAnf^Anc^ residue is <.3 in Nif^Anc^ and >.7 in VnfAnf^Anc^ and, for maintained sites, >.7 in Nif^Anc^ and >.7 in VnfAnf^Anc^.

We tracked the evolution of D-subunit residues through the emergence of the ancestral G-D interface. We identified 42 D-subunit sites interacting with the G-subunit following its initial recruitment in ancestral VnfAnf^Anc^ (**Supplementary Tables**). Of the 42 sites, 26 are well-conserved among extant Vnf/Anf nitrogenases and form a core interaction patch that was likely integral to the early G-D interface (**Figure 3E, 3F**). We inspected these 26 sites and found that nine sites evolved (i.e., were substituted) alongside G-subunit emergence, imposing subtle changes to the electrostatic surface potential of the VnfAnf^Anc^ interaction patch (**Figure S8, S9**).

Intriguingly, we find that multiple ancestral D-subunit surface features appear to have been preconditioned for G-subunit binding, collectively forming a “proto-interface” upon which interactions with the newly evolved G-subunit was built. Out of 26 residues within the core, 16 (∼62%) show that the D-subunit interaction patch had already become VnfAnf^Anc^-like prior to G-subunit emergence (**Figure 3F,G**). This residue-level conservation translates to maintained hydrophobicity (median apolar fraction ≈ .2) of the (proto-)interaction patch both before and after G-subunit recruitment (**Figure S10**). Further, shape complementarity as an integral feature of the G-D interface appears to have preceded the G-subunit. In *A. vinelandii* Vnf^21^, a cleft within the D-subunit interaction patch is occupied by a strictly conserved G-subunit residue, Leu18. This residue is among the most well-connected within the G-D interaction network (**Figure 3A-D**), is critical for interface stability (**Figure S7**), and is among the most allosterically active, capable of generating long-distance effects that reach across opposite ends of the nitrogenase complex (**Figure S11**). These effects reflect the importance of the G-D interface in controlling the global motions of the nitrogenase complex and evoke similar, long-distance structural perturbations that are induced through the nitrogenase catalytic cycle^56,57^. The cleft itself provides an important anchor point for G-subunit binding and is present in both ancestral and representative extant nitrogenase structures (**Figure 3A,B**, **Figure 4**).

**Figure 4.**
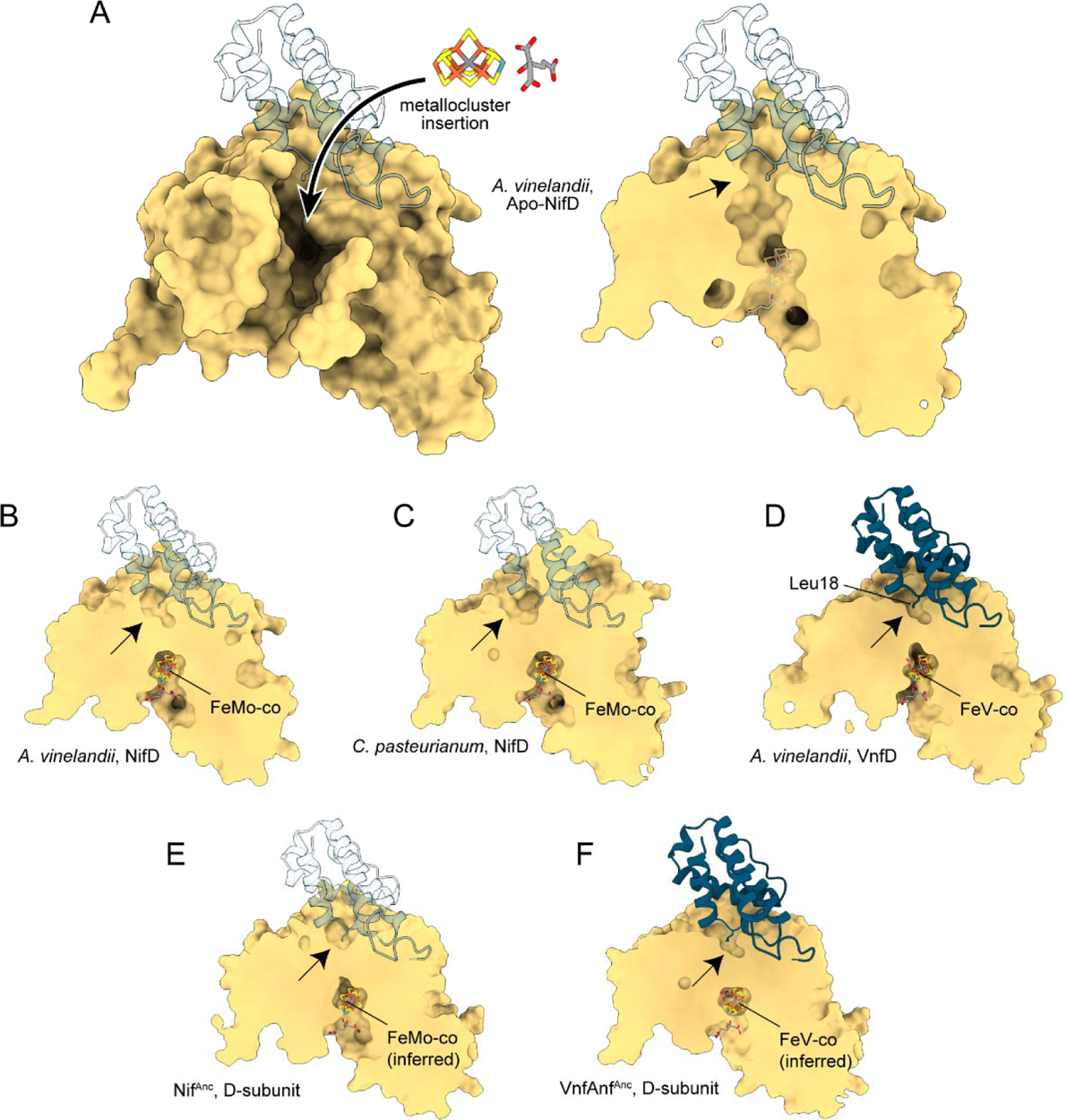
Conservation of the D-subunit cofactor insertion cleft across extant and ancestral nitrogenases. Small black arrow indicates position of the proposed cofactor insertion cleft. **A)** Crystallographic structure of the A. vinelandii apo-NifD protein (PDB 1L5H). Cross-section view (right) reveals the proposed metallocluster insertion site^58^, the entrance of which structurally aligns with the G-D interface of VnfD. Aligned, cross-section views are displayed for extant **B)** A. vinelandii NifD (PDB 3U7Q), **C)** Clostridium pasteurianum NifD (PDB 4WES), and **D)** A. vinelandii VnfD (PDB 5N6Y) crystallographic structures as well as predicted, ancestral **E)** Nif^Anc^ D-subunit and **F)** VnfAnf^Anc^ D-subunit structures. **(A-C, E)** A. vinelandii VnfG (PDB 5N6Y; transparent) is superimposed to enable comparison across nitrogenases with and without native G-subunit proteins. **(A-F)**

The functional significance of the D-subunit cleft provides an explanation for why the interface patch resisted significant remodeling with G-subunit addition, as well as implicates an adaptive role for the G-subunit. In extant Nif, the cleft aligns with the entrance to a proposed nitrogenase metallocluster insertion site, which, in the apo-NifD protein, adopts an open conformation to permit cluster loading^58^. Given the consistent presence of the cleft across Nif, Vnf, and Anf nitrogenases, a similar mechanism can be predicted for Vnf and Anf and is likely an ancient, universal feature of nitrogenase assembly that predates the G-subunit and was maintained due to its essentiality. In fact, we find that 12 interface residues surrounding the insertion site are well-conserved in extant Group III Nif clades that diverge prior to the origin of the G-subunit, and five of these are either universally or nearly universally conserved among all nitrogenases (**Figure S12**). Structural alignments indicate that the G-subunit caps the opening of the insertion site, with the side chain of the conserved Leu18 residue oriented into the cleft and toward the loaded active site. An unusual feature of the G-subunit, a carboxyl-terminal tail that terminates in a strictly conserved tyrosine residue (Tyr113), bridges the presumed opening of the apo-D-subunit protein and neighbors universally conserved D-subunit residues. The identified interactions between the bound G-subunit and the proposed metallocluster insertion site in the D-subunit implicate a role for the G-subunit in nitrogenase assembly, which is supported by experimental evidence that VnfG and VFe-co of *A. vinelandii* associate *in vitro*, as well as of the requirement of both VnfG and VFe-co to stabilize an active complex^23^. Conservation of G-subunit residues surrounding the cleft (and thus the metallocluster insertion site) across our phylogenetic dataset suggests that this putative role was early-evolved and maintained through its evolution.

### The G-subunit confers specialization between Vnf and Anf nitrogenases

We assessed to what degree sequence variation within the G-subunit is capable of driving the functional distinctions between Vnf and Anf isozymes, which are each dependent on distinct metalloclusters and exhibit varied N_2_-reduction efficiencies and reactivities to different chemical substrates^18,33-35^. We identified protein sites that are “specialized” between Vnf and Anf (i.e., diverge between Vnf and Anf but are otherwise conserved in each of the two groups), and therefore most likely drive functional variation. We find the degree of specialization (relative to protein size) within the G-subunit to be on par with that of the D-subunit but observe comparatively little specialization within the K-subunit (**Figure 5A, 5B, Supplementary Tables**). Therefore, the G-subunit contributes an outsized fraction of the total sequence specialization between Vnf and Anf, despite its remoteness from any nitrogenase metalloclusters. Specialized sites within the G-subunit are distal from the G-D interface. These include one residue close to the presumed H-subunit binding site, which might contribute to a previously proposed role in mediating H-subunit binding specificity^24^.

**Figure 5.**
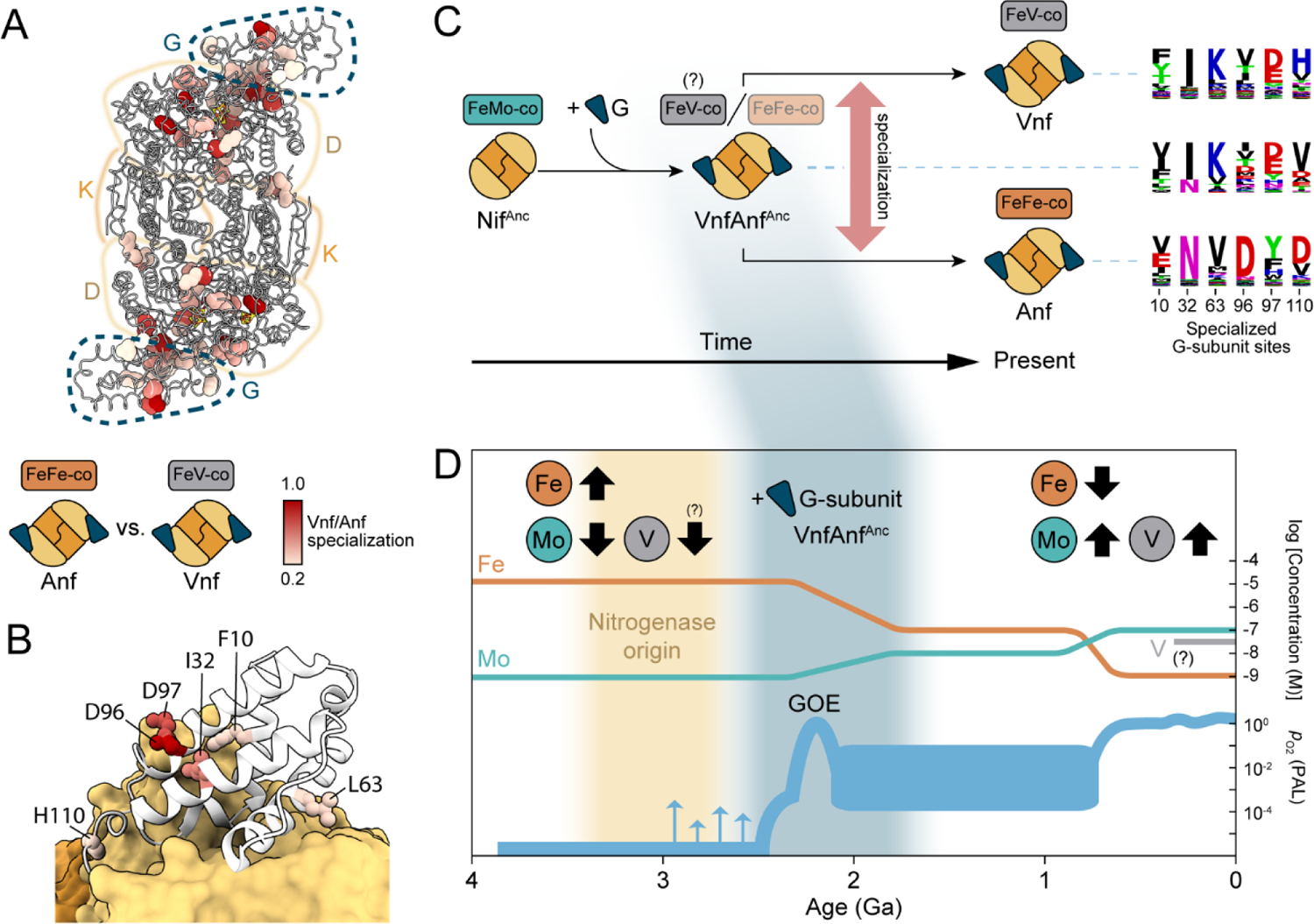
Coevolution of G-subunit specialization and Vnf/Anf metal dependence. **A)** Specialized sites mapped to the nitrogenase VnfDGK structure (PDB 5N6Y^21^). Scale in A) applies to B). **B)** Detailed view of specialized sites across the G-subunit. **C)** Evolution of specialized G-subunit sequence motifs through the divergence of Vnf and Anf. Sequence logos scale either with amino acid frequencies (for extant Vnf or Anf nitrogenases) or posterior probabilities (for ancestral VnfAnf^Anc^) at each site. VnfAnf^Anc^ dependence on FeV-co is inferred based on comparisons of specialized sequence motifs. **D)** The bulk marine concentrations of redox-sensitive metals involved in nitrogen fixation (Mo, V, Fe) have changed with progressive oxygenation of the Earth surface environment, with Fe more abundant in the Archean and Mo/V more abundant today. Though quantitative estimates of historical V concentrations are not presently available, oxygenation likely impacted the speciation and solubility of V and Mo similarly over geologic time^59,60^. Metal concentrations from Zerkle et al.^61^ and references therein and p_O2_ from Lyons et al.^27^. Age constraints of nitrogenase origins and VnfAnf^Anc^ (i.e., G-subunit emergence) from Stueken et al.^62^ and Parsons et al.^51^, respectively. GOE: Great Oxidation Event, PAL: present atmospheric level.

We next examined how these signatures of sequence-level specialization manifested early in G-subunit evolution, tracking their amino acid content through the divergence of Vnf and Anf clades. We find that the specialized sequence motifs of Vnf^Anc^ (Vnf ancestor) and Anf^Anc^ (Anf ancestor) most closely resemble those of their respective descendants (**Figure 5C**). The most likely Vnf^Anc^ motif across divergent sites is “FIKIDV” (extant Vnf consensus “FIKVDH”) whereas the most likely Anf^Anc^ motif is “VNVDYD” (extant Anf consensus “VNVDYD”). However, we find that the most likely motif of the common VnfAnf^Anc^ ancestor, “VIKIDV,” is more like that of Vnf than Anf, even accounting for the statistical uncertainty associated with ancestral reconstruction. This result suggests that the earliest G-subunit proteins were Vnf-like (in agreement with previous inferences of ancestral nitrogenases that indicate Vnf evolved prior to Anf^15^), and that the G-subunit contributed to specificity required for vanadium dependency early in the birth of a new nitrogenase isozyme.

## DISCUSSION

In the present study, we interrogated the origin, early evolution, and functional significance of a major novelty in the history of nitrogen fixation: the emergence of the G-subunit in Vnf and Anf nitrogenases >1.5 billion years ago^51^. With the birth of the nitrogenase G-subunit, not only did a rare evolutionary event occur with associated fitness costs overcome^4^, but a novel structural domain became fully integrated into the nitrogenase complex and was retained in one of the most central metabolic processes to Earth’s biosphere.

We demonstrate that there currently exists no candidate, evolutionary predecessor of the G-subunit family, whether another protein, protein fragment, or segment of non-genic DNA. This finding sets the origin of the G-subunit apart from other major evolutionary events in the history of nitrogen fixation that are explainable by gene duplication, including the divergence between other catalytic subunits, between nitrogenases and their maturation protein scaffolds, and between nitrogenase isozymes^32,63^. The addition of such a small and evolutionarily enigmatic protein domain appears to be an anomaly across biogeochemical enzymes.

We propose that the G-subunit was formed *de novo* from previously non-genic DNA, more recently acknowledged as a viable process for generating genetic novelty^47-49^. Such an event may have accompanied the genetic duplication and divergence of Nif that produced alternative nitrogenases^32^. Regardless of its originating mechanism, retainment of the G-subunit gene within the nitrogenase operon appears to have been improbable. Insertions between nitrogenase D- and K-subunit genes must either be rare or frequently deleterious, as evidenced by their small intergenic distance and absence of any other disrupting open reading frames across extant Nif clusters in our dataset. Thus, the evolution of the G-subunit constitutes an unprecedented gain in structural and genetic complexity in the evolutionary history of nitrogen fixation.

By analyzing predicted, nitrogenase ancestors, we reveal an evolutionary steppingstone in the initial recruitment of the G-subunit. Our reconstruction of ancestral, D-subunit surface features indicates that the G-subunit binding site was built from an essential, preexisting metallocluster insertion site, which additionally appears preconditioned for G-subunit interactions. Given that all known extant nitrogenases share this particular assembly scheme, its presence likely dates as far back as the oldest geochemical evidence of biological nitrogen fixation at ∼3.2 Ga^62^, hundreds of millions of years prior to the earliest whiffs of Earth oxygenation^52,53^. How then did this essential surface feature accommodate burial by a new interaction partner? Metallocluster delivery in extant nitrogenases is thought to be mediated by transient protein-protein interactions^58,64,65^ that would presumably have shaped nitrogenase surface properties, perhaps explaining the early evolved hydrophobicity of the ancestral, proto-interface. Our evolutionary analyses, coupled with experimental support of a role in nitrogenase assembly^23^, suggest that the G-subunit was integrated within the assembly pathway, replacing and/or mediating earlier evolved protein or metallocluster interactions. Thus, the birth of the permanent G-D interface represents a gradual subsummation of formerly transient interactions that provided an evolutionary “cradle” for a permanent, novel structural domain.

In its capacity to mediate newly evolved incorporation of V- and Fe-based clusters, the G-subunit would have enabled the evolutionary burst of both metal dependence and ecological diversity within the Group III nitrogenase lineage from which alternative nitrogenases originated. We find that G-subunit residues, despite being relatively distal from the active-site and not permanently retaining bound, reactive metalloclusters like other nitrogenase subunits, have an outsized contribution to the sequence-level specialization between alternative nitrogenases. Possible functional outcomes include improved specificity toward metalloclusters or associated protein partners during nitrogenase assembly (preventing costly mismetallation^66^)^23^ and mediating crosstalk between electron transfer and catalytic nitrogenase components^24^. Each of these functions would have equipped diazotrophs to coordinate multiple nitrogenases systems with unique metal dependencies within the same cellular environment. This ability is advantageous in ancient or modern environments that experience transient molybdenum limitation, with alternative nitrogenases providing a “fail-safe” strategy under conditions when molybdenum-dependent nitrogen fixation cannot be supported^67^. The G-subunit therefore primed these organisms for new, molybdenum-limited ecological niches, as well as for fluctuations in redox-sensitive metal availabilities following the rise of oxygenic phototrophs^27,68^, one of the most significant ecological revolutions in Earth history.

An ongoing debate in evolutionary biology centers on the degree to which environmental conditions drive biological innovations. In the evolution of biological nitrogen fixation, an intuitive narrative is that the diversification of nitrogenase metal dependence, intimately tied to the productivity and ecological distribution of nitrogen fixation, was triggered largely by transitions in bulk, environmental metal availabilities beginning ∼2.5 to 3.0 Ga^26,52,53^, the historical records of which are primarily built from geochemical analyses of ancient marine sediments (**Figure 5D**). Our study highlights the additional importance of molecular evolutionary events emerging from specific historical trajectories that predate, and are therefore removed from, candidate environmental triggers. The birth of the nitrogenase G-subunit, by providing a crucial component in novel metal incorporation, expanded the adaptive landscape for biological nitrogen fixation. This ancient, orphan protein was birthed by seemingly improbable events but its recruitment was nevertheless enabled by a conserved, nitrogenase assembly pathway that arose hundreds of millions of years prior to the rise of Earth surface oxygen and the accompanying shifts in metal availabilities. Whether this evolutionary transition in the history of biological nitrogen fixation could have been crossed in the absence of the G-subunit awaits further studies. Similar, future synergies between historical studies of molecular novelties and the environmental circumstances that shaped them can refine our understanding of how the Earth-life planetary system has evolved over billions of years.

## METHODS

### Nitrogenase protein sequence curation

A preliminary 412-sequence dataset of nitrogenase Vnf/AnfG proteins was compiled by BLASTP^39^ search (Expect value threshold = 0.1) against the National Center for Biotechnology Information (NCBI) non-redundant protein sequence database (nr)^69^ (accessed January 2022), using the *A. vinelandii* VnfG query sequence (WP_012698949.1). BLASTP hits were curated by filtering against partial sequences and homologs whose encoding genes are distantly located from *vnf* or *anf* operons (and thus unlikely to be canonical *vnf/anfG* genes). Certain VnfG sequences were manually extracted from fused VnfDG proteins, using MAFFT v7.490^70^ alignments to other D- or G-subunit sequences as a guide. The preliminary Vnf/AnfG sequence dataset was pruned by CD-HIT^71^ (sequence identity threshold = 90%), yielding a final 188-sequence G-subunit dataset.

### G-subunit homolog search

In addition to the BLASTP search described above, three sequence-based search tools– TBLASTN^39^, PSI-BLAST^40^, and HMMER^41^–were used to identify distantly related homologs of the nitrogenase G-subunit (conducted January 2022). TBLASTN and PSI-BLAST searches against the NCBI nucleotide collection (nr/nt) and nr protein database, respectively, were executed with the *A. vinelandii* VnfG query (WP_012698949.1), an Expect value threshold = 0.1, and word size = 2 (all other parameters default). HMMER search (hmmsearch) against the UniProt Reference Proteome database^72^ was performed using an HMM profile generated from our MAFFT-aligned, 188-sequence G-subunit dataset (see above) with an Expect value threshold = 0.1 for both sequences and hits (all other parameters default).

Structure-based searches of distant G-subunit homologs were performed by the FoldSeek server^42^ using the *A. vinelandii* VnfG crystallographic structure (PDB 5N6Y^21^) query against the Protein Data Bank and AlphaFold-UniProt/Swiss-Prot databases. We employed both 3Di-AA and TM-Align modes with an 3Di-AA Expect value threshold = 0.01 or a TM-score = 0.5 (template modeling score).

### Phylogenetic and ancestral sequence reconstruction

The curated, 188-sequence Vnf/AnfG dataset was aligned by MAFFT. No outgroup to G-subunit sequences was included because the distant homology search described above did not yield convincing hits (see **Results**). IQ-TREE v1.6.12^73^ was used to reconstruct a maximum-likelihood G-subunit protein phylogeny, using the best-fit evolutionary model, WAG+R6 (selected by the Bayesian Information Criterion, BIC), tested by ModelFinder^74^, with branch support calculated by ultra-fast bootstrap approximation (UFBoot)^75^ and the Shimodaira-Hasegawa-like approximate likelihood-ratio test (SH-like aLRT)^76^.

A phylogeny of concatenated Vnf/AnfHDGK proteins was generated by retrieving syntenous subunit sequences to the curated VnfG/AnfG dataset. Certain VnfH sequences that recently diverged from NifH clades were omitted due to having different evolutionary histories than those of other Vnf/Anf proteins. NifHDK sequences from a prior phylogenetic analysis^77^ were included as an outgroup (n = 67, closely branching to Anf/Vnf) and were used to calculate intergenic *nifD*-to-*nifK* distances (n = 306). Each subunit sequence dataset was aligned by MAFFT prior to concatenation. Maximum-likelihood phylogenetic reconstruction was performed by IQ-TREE as described above (best-fit evolutionary model, BIC: LG+F+R9).

Marginal maximum-likelihood ancestral sequence reconstruction of concatenated Vnf/AnfHDGK sequences was performed by PAML v4.9j^78^ (LG+F+G4 model). Mean sitewise posterior probabilities D- and G-subunit ancestors analyzed in this study range from .75 to .96 (**Table S1**). Sequence gaps were reconstructed as described by Aadland et al.^79^. Briefly, the input protein sequence alignment was recoded as a binary amino-acid presence/absence matrix. Ancestral presence/absence states were reconstructed by PAML using a binary character model and state frequencies inferred by maximum likelihood.

Logo representations of extant multiple sequence alignments were generated with the Python library LogoMaker^80^ using the probability transformation of the count matrix. Representations of ancestral sequences were generated from the posterior probability matrices produced by PAML. Sequence and phylogenetic tree datasets are available at https://github.com/kacarlab/nitrogenase-g-subunit.

### Optimal growth temperature prediction

For taxa represented in the nitrogenase G-subunit sequence dataset, optimal growth temperatures (OGT) were predicted from genomic content using the method described by Sauer and Wang^81^. Due to genome incompleteness and the absence of 16S rRNA sequences in fragmented genomes for certain representative taxa, the selected “Superkingdom Bacteria” and “Superkingdom Archaea” regression models for OGT prediction excluded genome size and 16S rRNA.

### Protein conservation and divergence analysis

Sequence conservation for D- and G-subunit proteins was determined by the ConSurf server^82^. Sites were considered well-conserved if their ConSurf grade were >7. Sequence specialization was determined by the TwinCons package^83^. TwinCons was run separately on individual alignments of extant H, D, G, and K subunit protein sequences with user-defined Vnf and Anf groups, using default parameters and the LG substitution matrix. TwinCons specialization scores were normalized to a 0-to-1 scale per analyzed subunit.

### Structural Modeling

Ancestral and extant NifHDGK protein structures were predicted using ColabFold^84^, which combines the AlphaFold2 method for structure prediction from multiple-sequence alignments^18^ with the MMSeq2 method to obtain evolutionary information^85^. Specifically, the colabfold_batch v1.4.0 implementation (https://github.com/YoshitakaMo/localcolabfold<u>) was used with</u> the AlphaFold-v2.0-multimer prediction model and standard options (3 recycles and AMBER all-atom optimization in GPU).Protein structures were visualized by ChimeraX^86^.

### Structure Analysis

Residue interaction networks^87^ were generated using ProLif^88^, which finds the different possible interactions (hydrophobic, electrostatic, hydrogen bonds, etc) between two groups of atoms (here defined as G- and D-subunit groups) within a structure snapshot (e.g. single structure, trajectory frame). Interactions were compiled into networks by NetworkX^89^, which were later visualized in Gephi^90^. Scripts used to generate the networks are available at https://github.com/kacarlab/nitrogenase-g-subunit.

Surface electrostatic maps were computed using the Poisson Boltzmann Solvent-Accessible model implemented in APBS^91^ through the PyMOL plugin (Schrödinger, Inc), using standard PDB2PQR and APBS protocols at pH 7.

Sitewise contribution to D-G interface stability was computed using *in-silico* alanine scanning, which substitutes residues with a minimal methyl moiety (alanine)^92^. The changes on the binding free energy values upon substitution were obtained using the machine-learning enabled mCSM2 method^93^ with default parameters.

Allosteric effects across the nitrogenase complex was modeled using Perturbation Response Scanning^94^, implemented in ProDy^94^ with default parameters and provided with PDB 5N6Y^21^ alpha carbon coordinates. Scripts used to model allosteric effects are available at https://github.com/kacarlab/nitrogenase-g-subunit.

## ACKNOWLEDGEMENTS

This research was supported by the National Aeronautics and Space Administration (NASA) Interdisciplinary Consortium for Astrobiology Research: Metal Utilization and Selection Across Eons, MUSE (19-ICAR19_2–0007), the NASA Arizona Space Grant (B.M.C), the John Templeton Foundation (B.K.; 61926), the National Science Foundation (B.K.; 2228495), the NASA Early Career Faculty Award (B.K.), the Hypothesis Fund Award (B.K.), and the Universidad Politécnica de Madrid (UPM) Margarita Salas Fellowship, founded by the Unión Europea - Next Generation EU (B.C.-Z.; UP2021-035). We thank Morgan Sobol for assistance with optimal growth temperature prediction, as well as Ariel Anbar, Alex Rivier, Derek Harris, Holly Rucker, Joanna Masel, Lance Seefeldt and Steven Russell for valuable discussions and feedback.

