## Supplemental Information for "The nitrogenase G-subunit is an ancient orphan protein that drove the ecological expansion of nitrogen fixation"

### Supplementary Information

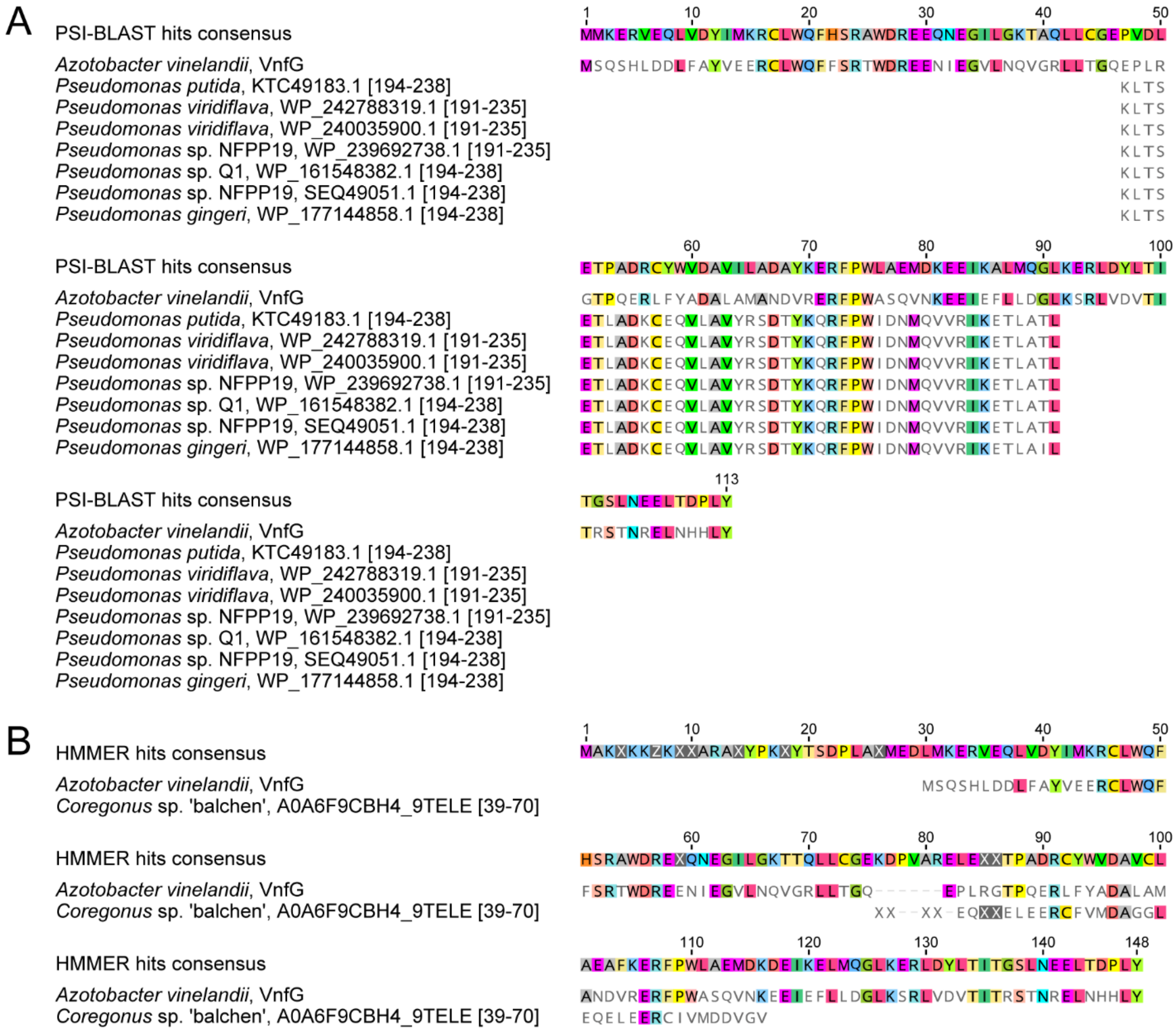


**Figure S1. Protein multiple sequence alignments of distant homology search hits to canonical G-subunit sequences. (A)** PSI-BLAST search results. **(B)** HMMER search results. **(A-B)** Only aligned regions of search hits are displayed, with site indices from the original sequences noted in brackets beside the sequence source organism and accession.

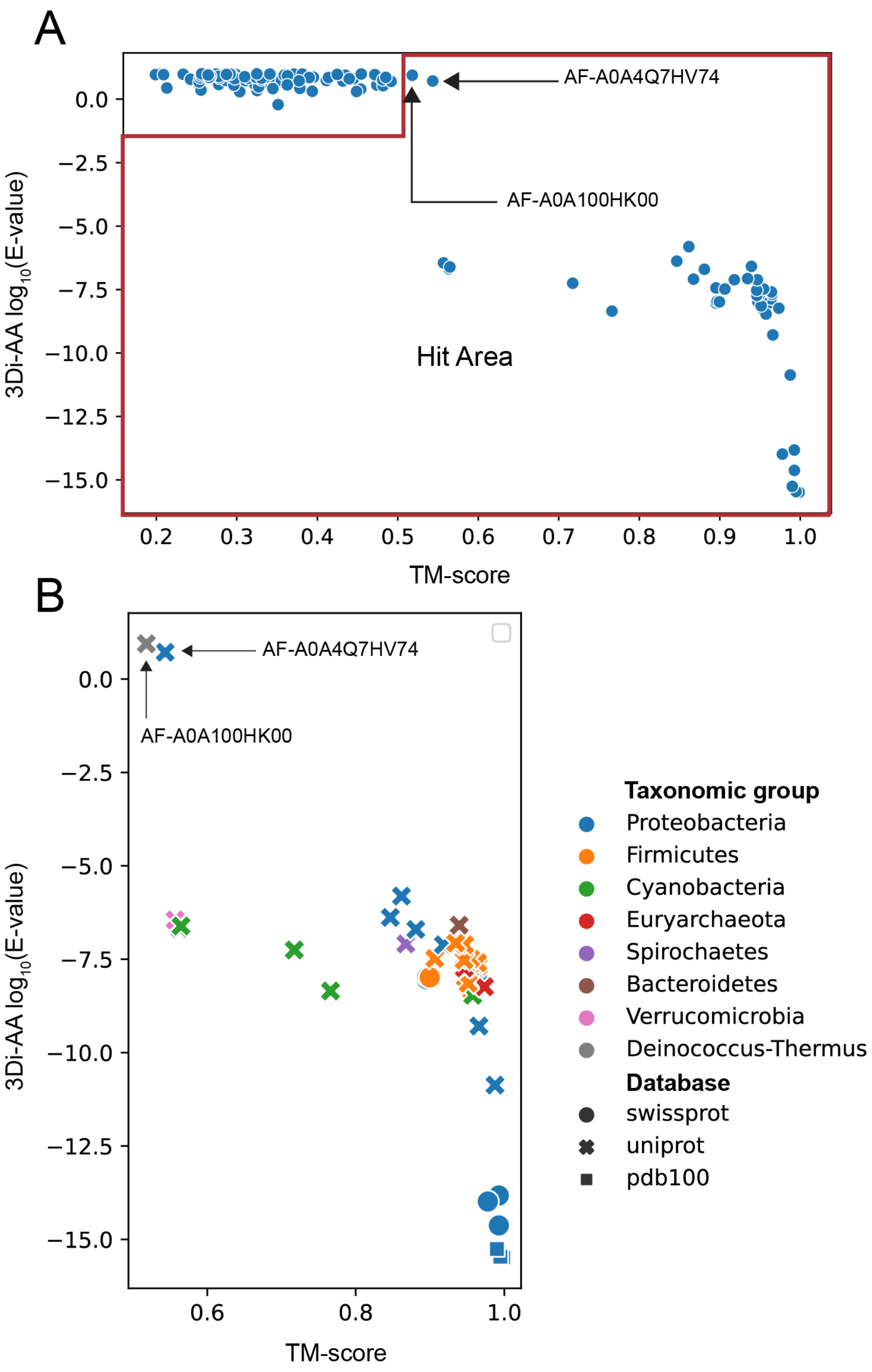


**Figure S2. 3Di-AA and TM-score distributions across A) all FoldSeek results and B) significant FoldSeek hits (3Di-AA $\log_{10}$(Expect value) < -2 or TM-score > 0.5).** Distant hits that are not canonical G-subunit sequences are labeled.

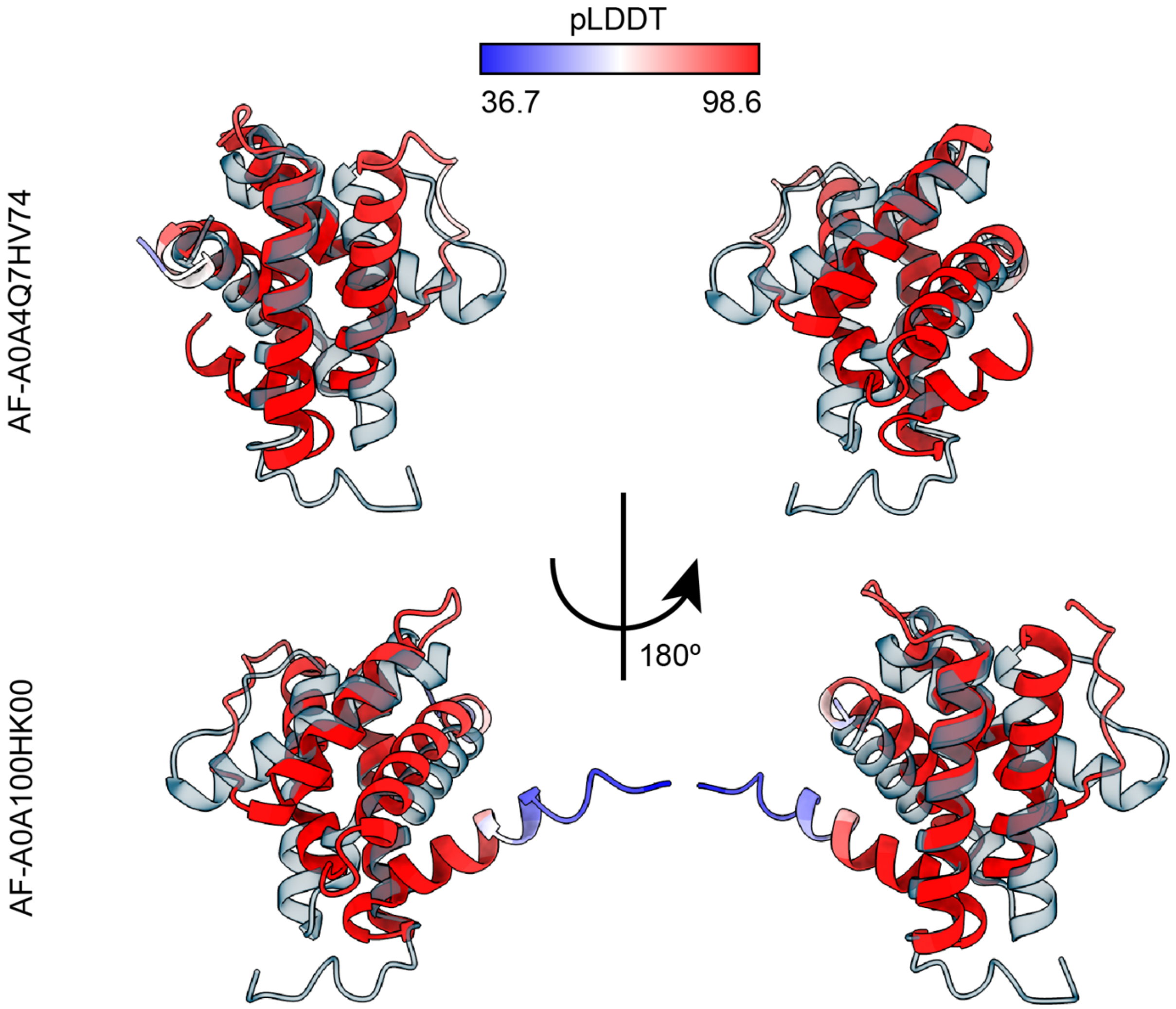


**Figure S3. Structural alignments between FoldSeek search query (VnfG, PDB 5N6Y[1]; transparent) and significant hits (3Di-AA $\log_{10}$(Expect value) < -2 or TM-score > 0.5). (A)** Uncharacterized protein (AlphaFold Database accession A0A100HK00) from *Deinococcus grandis* (3Di-AA $\log_{10}$(Expect value) = 0.94, TM-score = 0.518). **(B)** Uncharacterized protein (AlphaFold Database accession A0A4Q7HV74) from *Vibrio vulnificus* (3Di-AA $\log_{10}$(Expect value) = 0.72, TM-score = 0.543). (A-B) FoldSeek hits colored by AlphaFold confidence metric (pLDDT), where a higher pLDDT denotes higher confidence in the predicted structural region.

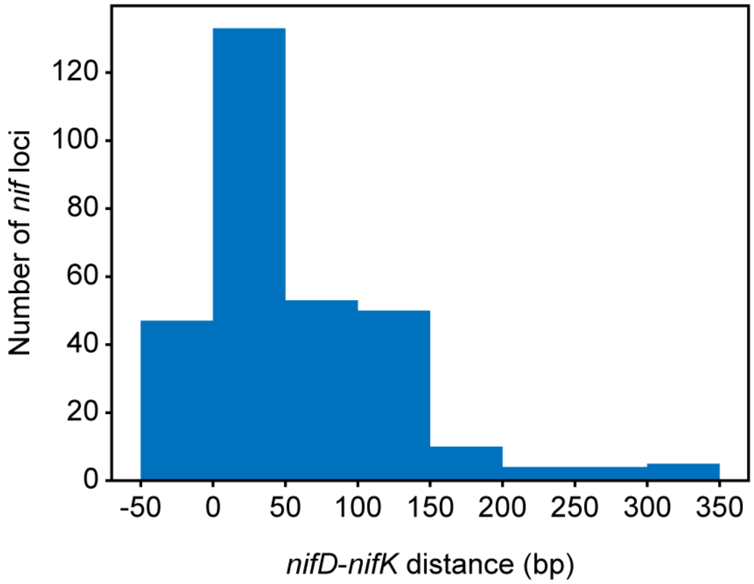


**Figure S4. Distribution of intergenic *nifD*-to-*nifK* distances across taxonomically diverse *nif* loci.** Negative values indicate overlapping *nifD* and *nifK* genes.

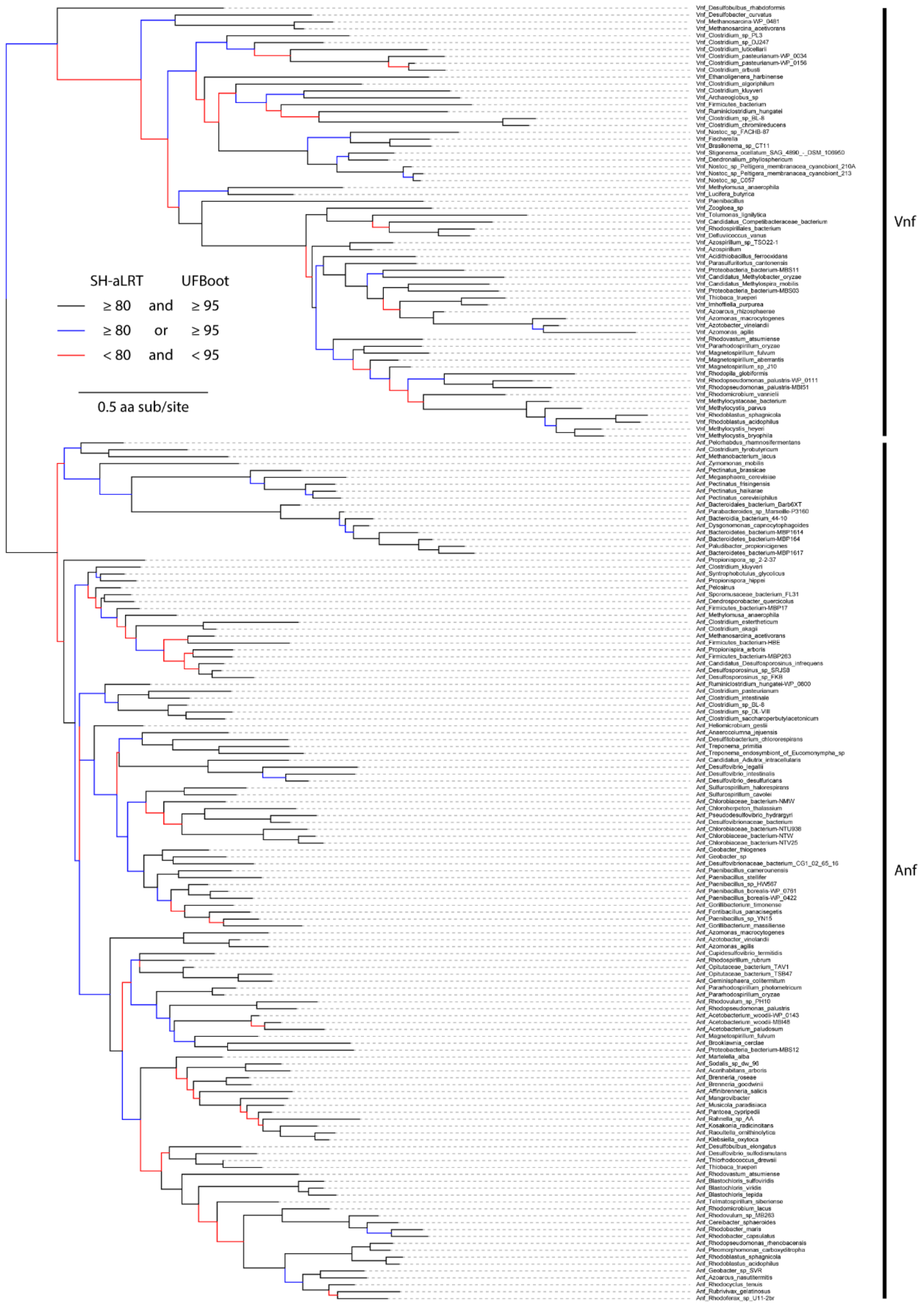


**Figure S5. Maximum-likelihood phylogeny of Vnf/AnfG protein sequences.** SH-aLRT: Shimodaira-Hasegawa-like approximate likelihood-ratio test, UFBoot: Ultrafast boostrap approiximation.

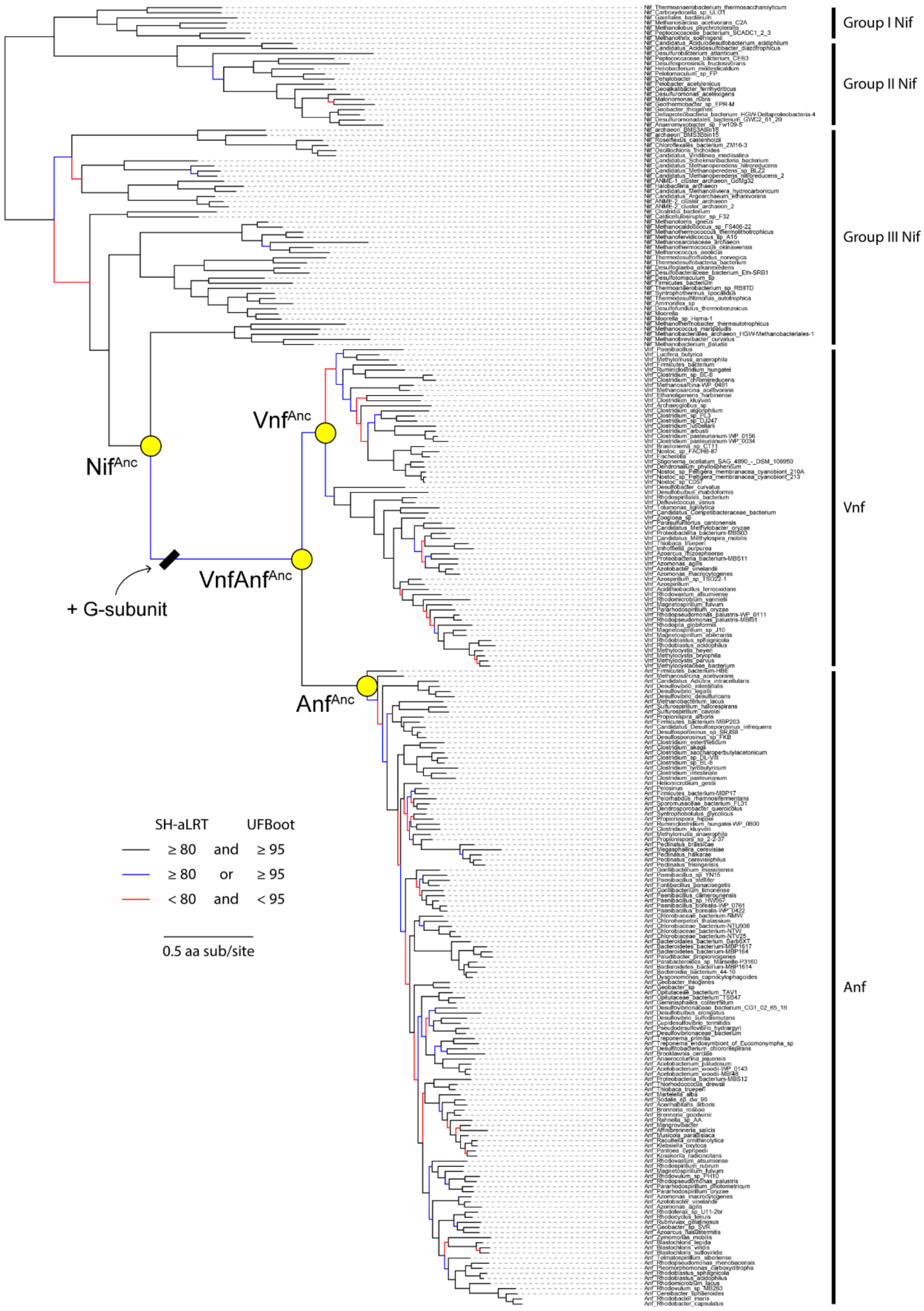


**Figure S6. Maximum-likelihood phylogeny of concatenated Nif/Vnf/AnfHDGK protein sequences.** SH-aLRT: Shimodaira-Hasegawa-like approximate likelihood-ratio test, UFBoot: Ultrafast boostrap approixmation.

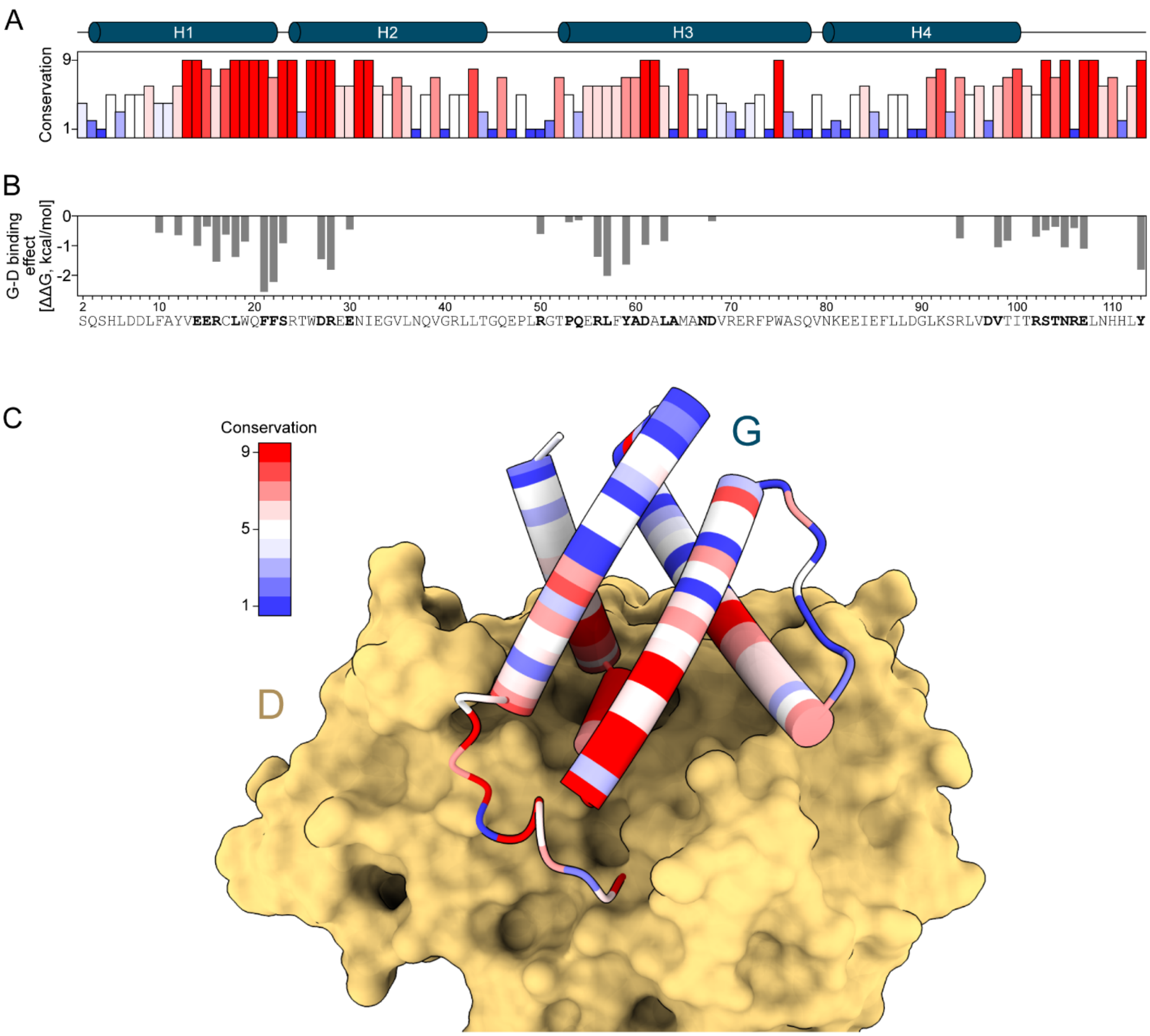


**Figure S7. G-subunit site conservation correlates with proximity to the G-D interface. (A)** Sitewise conservation scores (from 1, low, to 9, high) across Vnf/AnfG proteins. G-subunit secondary structure indicated at top. **(B)** Sitewise contribution to G-D interface stability, assessed by *in silico* alanine scanning, where a lower ΔΔG indicates greater decrease in G-D binding energy upon alanine substitution. *A. vinelandii* sequence shown at bottom also applies to (A) Bold amino acids are within the predicted G-D residue interaction network (see **Materials and Methods**).

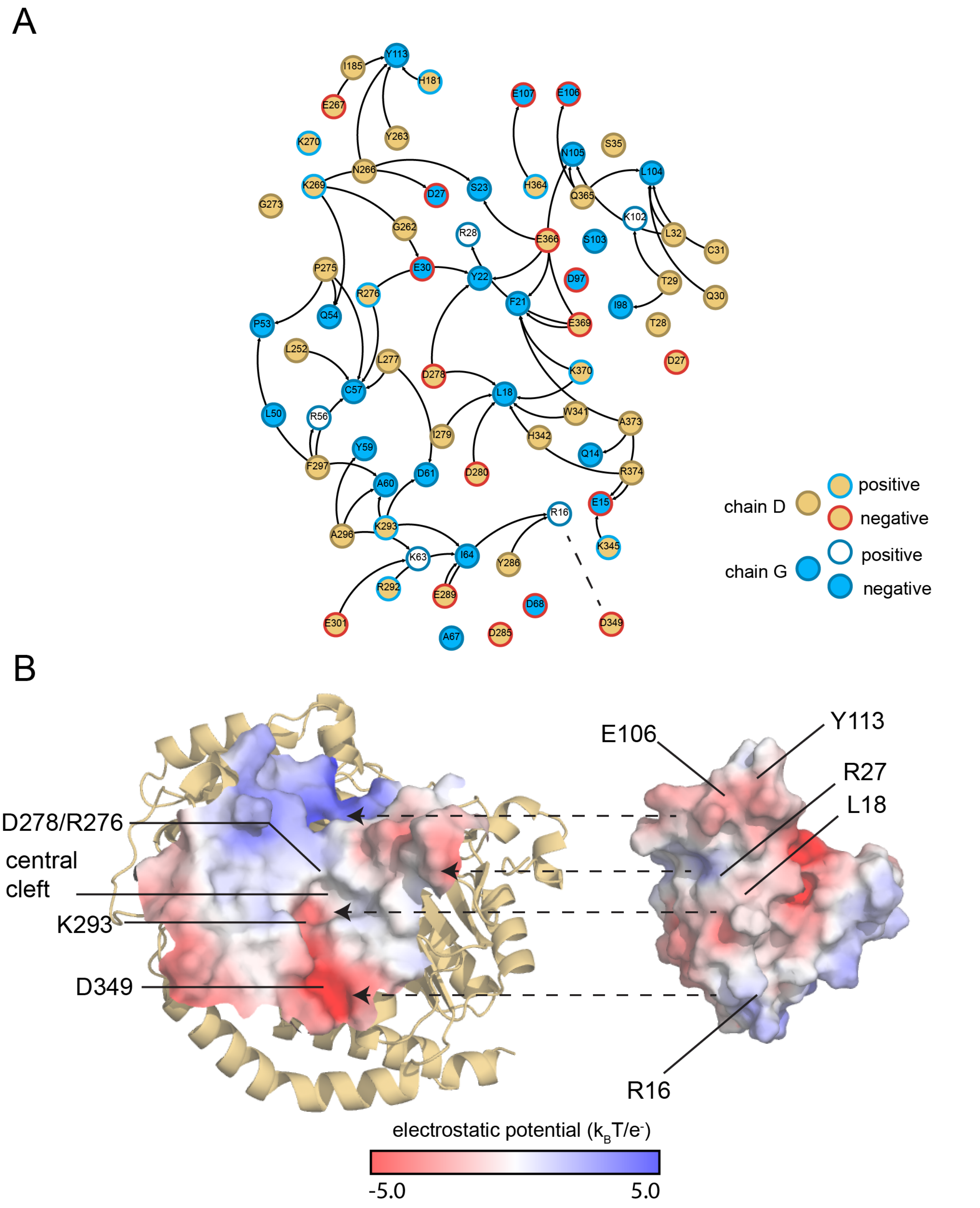


**Figure S8. Electrostatic surface potential of the ancestral VnfAnf$^{Anc}$ G-D interface.** **(A)** VnfAnf$^{Anc}$ residue interaction network. Nodes are colored by subunit (fill) and charge (outline). **(B)** Electrostatic complementarity of D- (left) and G-subunit (right) VnfAnf$^{Anc}$ proteins.

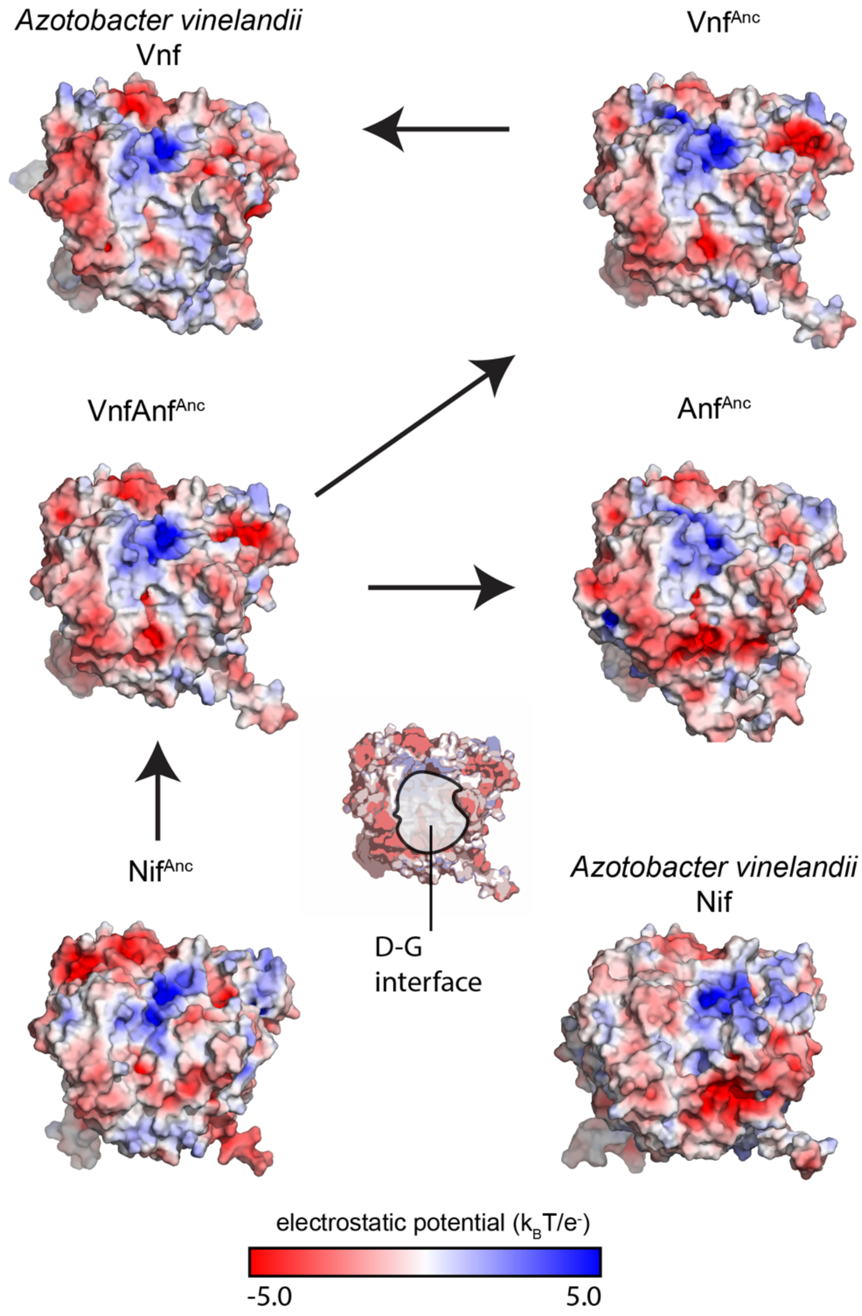


**Figure S9. Electrostatic profiles of extant and ancestral D-subunit proteins.** The central cartoon shows the location of the D-G interface patch.

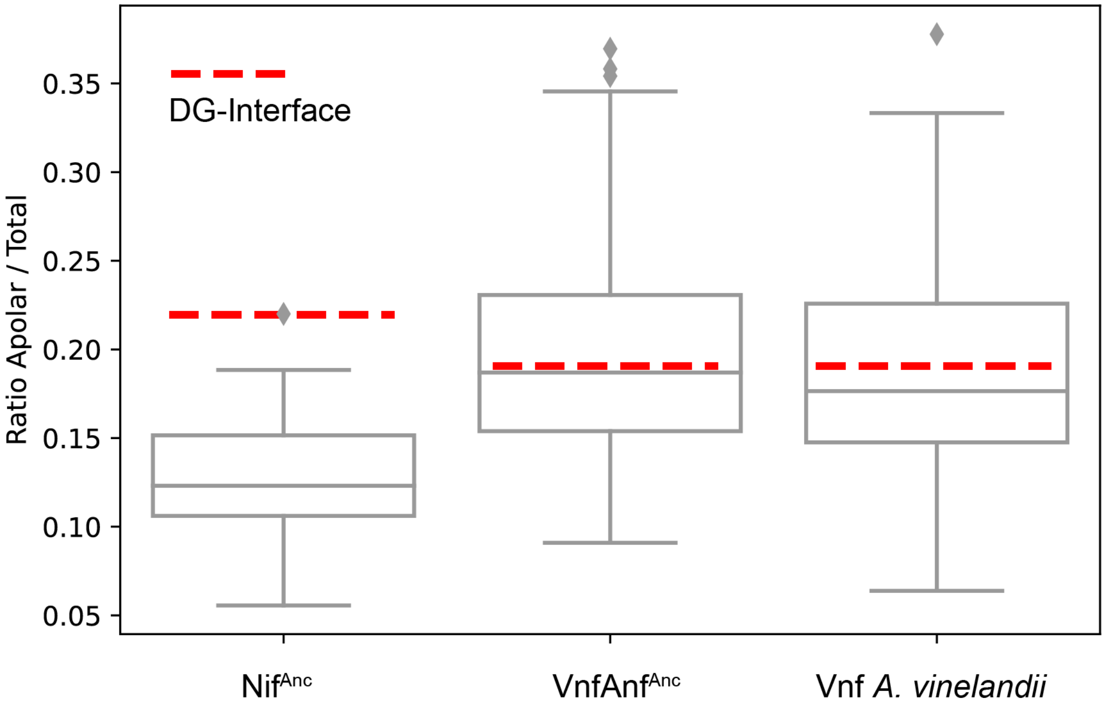


**Figure S10. Hydrophobicity of ancestral and extant D-subunit residues within the G-D interface.** The hydrophobic fraction of the D-G interface (or proto-interface) is shown by a dashed red line. Boxplots represent the hydrophobic fraction distribution of sampled surface patches containing D-subunit residues, excluding all protein-protein interface residues.

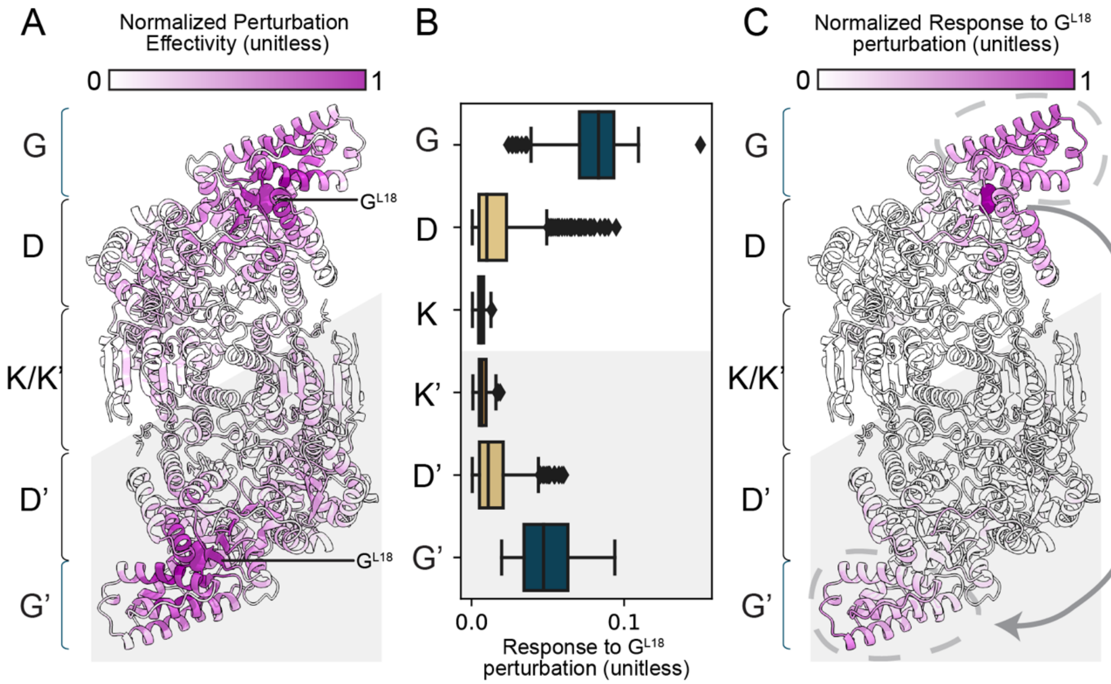


**Figure S11. Allosteric signaling generated by the nitrogenase G subunit. A)** Effectiveness of allosteric signaling mapped to Vnf (PDB 5N6Y), as calculated by perturbation response scanning and normalized on a 0-to-1 scale. **B)** Sensitivity to allosteric signaling after perturbation of G-Leu18. Box plots represent perturbation responses calculated across all residues in each protein subunit. **C)** Sensitivity in B) mapped to Vnf (PDB 5N6Y), normalized on a 0-to-1 scale.

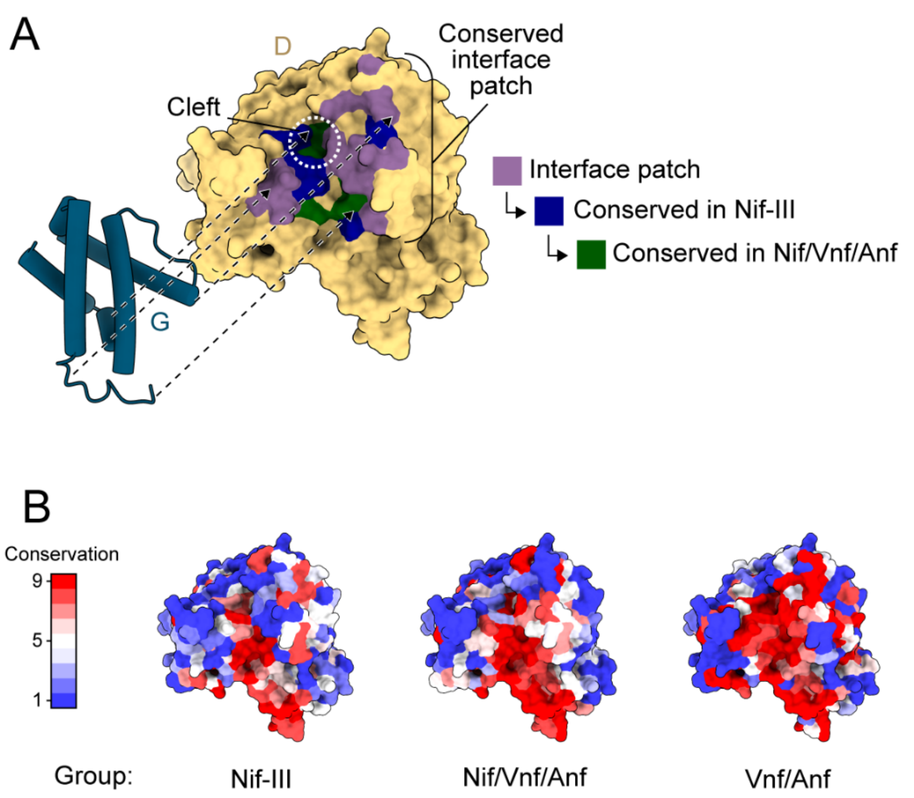


**Figure S12. Conservation of the D-subunit interface patch across different nitrogenase clades. A)** D-subunit interface patch (PDB 5N6Y), colored by nested criteria: 1) located within the interface patch (purple), 2) conserved among Group III Nif ("Nif-III"; dark blue), and 3) conserved among all nitrogenases (Nif/Vnf/Anf; green). **B)** Conservation of the D-subunit service in Group III Nif, all nitrogenases (Nif/Vnf/Anf), or alternative nitrogenases (Vnf/Anf). All structures are oriented similarly to A).

### Table S1. Statistical support for ancestral sequences analyzed in this study.

| Node | Protein | Length (aa) | Mean sitewise posterior probability (± 1 SD) | Percentage ambiguous sites (posterior probability < 0.7) |
|---|---|---|---|---|
| $Nif^{Anc}$ | D-subunit | 456 | 0.88 ± 0.19 | 18% |
| | G-subunit | n/a | n/a | n/a |
| $VnfAnf^{Anc}$ | D-subunit | 471 | 0.90 ± 0.17 | 14% |
| | G-subunit | 112 | 0.75 ± 0.24 | 39% |
| $Vnf^{Anc}$ | D-subunit | 471 | 0.96 ± 0.11 | 5% |
| | G-subunit | 112 | 0.85 ± 0.20 | 22% |
| $Anf^{Anc}$ | D-subunit | 519 | 0.95 ± 0.13 | 8% |
| | G-subunit | 116 | 0.90 ± 0.18 | 13% |
